# The epigenetic evolution of gliomas is determined by their IDH1 mutation status and treatment regimen

**DOI:** 10.1101/2021.08.09.455687

**Authors:** Tathiane M Malta, Thais S Sabedot, Indrani Datta, Luciano Garofano, Wies Vallentgoed, Frederick S Varn, Kenneth Aldape, Fulvio D’Angelo, Spyridon Bakas, Jill S Barnholtz-Sloan, Hui K Gan, Mohammad Hasanain, Ann-Christin Hau, Kevin C Johnson, Mustafa Khasraw, Emre Kocakavuk, Mathilde CM Kouwenhoven, Simona Migliozzi, Simone P Niclou, Johanna M Niers, D. Ryan Ormond, Sun Ha Paek, Guido Reifenberger, Peter A Sillevis Smitt, Marion Smits, Lucy F Stead, Martin J van den Bent, Erwin G Van Meir, Annemiek Walenkamp, Tobias Weiss, Michael Weller, Bart A Westerman, Bauke Ylstra, Pieter Wesseling, Anna Lasorella, Pim J French, Laila M Poisson, The GLASS Consortium, Roel GW Verhaak, Antonio Iavarone, Houtan Noushmehr

## Abstract

Tumor adaptation or selection is thought to underlie therapy resistance of gliomas. To investigate the longitudinal epigenetic evolution of gliomas in response to therapeutic pressure, we performed an epigenomic analysis of 143 matched initial and recurrent patients with IDH-wildtype (IDHwt) and IDH-mutant (IDHmut) gliomas. IDHwt gliomas showed a longitudinally stable epigenome with relatively low levels of global methylation, whereas the epigenome of IDHmut gliomas showed initial high levels genome-wide of DNA methylation that was progressively reduced to levels similar to those of IDHwt tumors. By integrating DNA methylation and gene expression data, adaptive changes of putative master regulators of the cell cycle and of differentiation were seen in IDHmut recurrent tumors. Furthermore, relapses of IDHmut tumors were accompanied by histological progression which in turn influenced survival, as validated in an independent cohort. Finally, the initial cell composition of the tumor microenvironment differed between IDHwt and IDHmut tumors and changed differentially following treatment, suggesting increased neo-angiogenesis and T-cell infiltration upon treatment for IDHmut gliomas. Our study provides one of the largest cohorts of paired glioma samples profiled with epigenomics, transcriptomics and genomics; and our results demonstrate that the treatment of IDHmut gliomas reshapes the epigenome towards an IDHwt-like phenotype. Accordingly, the prevalent practice of early genotoxic treatment in this patient population may need to be revisited.

## Introduction

Despite advances in our biological understanding, molecular classification and surgical techniques, management of diffuse gliomas of adulthood remains challenging making it an incurable disease (Malta et al., 2018a). Compared to gliomas of the same grade that carry intact isocitrate dehydrogenase (*IDH*) 1 and 2 genes, gliomas with IDH mutations exhibit a less aggressive clinical course which has led to their separation as distinct tumor types in the 2016 World Health Organization (WHO) classification of Tumors of the Central Nervous System (CNS) (Louis et al., 2016). Based on the revised 2021 WHO Classification (Louis et al., 2021), IDH-mutant tumors now comprise two distinct tumor types, namely “oligodendroglioma, IDH-mutant and 1p/19q-codeleted CNS WHO grade 2 or 3”, and “astrocytoma, IDH-mutant CNS WHO grade 2, 3 or 4”. Yet, there is controversy on the morphological criteria used to distinguish CNS WHO grades 2 and 3, and homozygous *CDKN2A* loss (a signature lesion of CNS WHO grade 4 among IDH-mutant astrocytomas), which is presently the only diagnostic molecular marker in these tumors (Shirahata et al., 2018; Brat et al., 2020; Tesileanu et al., 2021). Thus, additional molecular characterization is needed to establish which of these tumors will rapidly progress or remain quiescent for several years with or without adequate therapy (Weller et al., 2021).

Epigenetics play a vital role in stratifying CNS tumors and gliomas into clinically relevant subtypes (Capper et al., 2018; Ceccarelli et al., 2016). The Glioma-CpG Island Methylator Phenotype (G-CIMP) discovered in IDH-mutant gliomas in 2010 (Noushmehr et al., 2010) provided an explanation for their favorable prognosis. Further investigation revealed a subset of IDH-mutant gliomas that presented with a lower degree of DNA methylation and poorer outcome, named G-CIMP-low; distinct from the previously described highly methylated tumors that have a better outcome, and renamed as G-CIMP-high (Ceccarelli et al., 2016). In our previous study, longitudinal epigenomics analyses of 77 paired glioma samples revealed that 12% of G-CIMP-high cases progressed to G-CIMP-low cases (de Souza et al., 2018) highlighting the importance of further studying epigenetic reprogramming following treatment and integrating it with other genomic platforms to discover new prognostic markers and therapeutic targets. Longitudinal analyses that integrate epigenetic and transcriptomic data offer an opportunity to define key master regulators driving the malignant progression of each glioma entity characterizing these prognostic subtypes as tumors transition to recurrence after initial clinical management.

Current management and treatment for gliomas include surgery followed by radiotherapy and/or alkylating chemotherapy (e.g. temozolomide [TMZ]). Recent studies have revealed fundamental molecular genetic changes associated with glioma treatment including the development of a hypermutation phenotype (Johnson et al., 2014; Hunter et al., 2006), increase in small deletion burden and acquisition of *CDKN2A* homozygous deletions associated with radiotherapy and acquired aneuploidy associated with cell cycle related genes and overall poorer outcome (Barthel et al., 2019; Kocakavuk et al., 2021). Interestingly, not all TMZ gliomas develop a hypermutator status which challenges the possible mechanisms driving this TMZ treatment-induced molecular phenotype (Mathur et al., 2020; Touat et al., 2020). However, treatment-induced hypermutation related to epigenetic and transcriptomic changes has not been well explored. It offers opportunities to identify potential epigenetic markers for predicting transition to a hypermutator status.

In the current study, by leveraging the Glioma Longitudinal AnalySiS (GLASS) international consortium (Barthel et al., 2019; GLASS Consortium, 2018; Varn et al., 2021), we analyze an epigenetic cohort of 143 glioma patients with matched initial and recurrent tumors, and include additional molecular data and clinical data to characterize the evolution of both IDH-wildtype and IDH-mutant gliomas. This is the largest cohort of paired initial and recurrent glioma samples profiled with epigenomics, transcriptomics and genomics, that we know of being used in the literature. Also, the GLASS-NL (GLASS in the Netherlands), a collaboration of several centers in the Netherlands treating patients with glioma, was included in this study to evaluate the effects of treatment in the epigenome of gliomas in an independent cohort. This consortium has collected material from 100 IDH-mutant astrocytoma patients who underwent at least two surgical resections (surgical interval > 6 months). Our study aimed at identifying key master regulators of tumor progression, identifying changes in the tumor microenvironment and epigenetic drivers of glioma evasion to treatment and examining differences in these processes between IDH-wildtype and IDH-mutant gliomas to derive better informed tailored treatments.

## Results

### Molecular evolution of matched initial and recurrent gliomas

The GLASS-international DNA methylation cohort consists of 143 patients with high-quality molecular data from at least two-time points, resulting in a total of 357 samples profiled by either Illumina 450K or EPIC Beadchip methylation arrays (Tables S1 and S2). The patients at initial diagnosis represented the three major glioma subtypes defined by DNA methylation signatures: IDH-mutant and 1p/19q-co-deleted oligodendroglioma (IDHmut-codel; n = 14); IDH-mutant astrocytoma without 1p/19q co-deletion (IDHmut-noncodel; n = 63); and IDH-wildtype glioblastoma (IDHwt; n = 66). We did not identify samples with IDH2 mutation and thus IDH here represents IDH1 only. Among the 143 patients with profiled DNA methylation, 59 patients had RNA sequencing data, 69 had DNA sequencing genomic data, either whole genome sequencing (WGS) or whole exome sequencing (WXS), and 54 had all three molecular data sets (Figures 1A and S1A). We selected two time-separated tumor samples for each patient, termed initial and recurrence, for further analysis.

**Figure 1.**
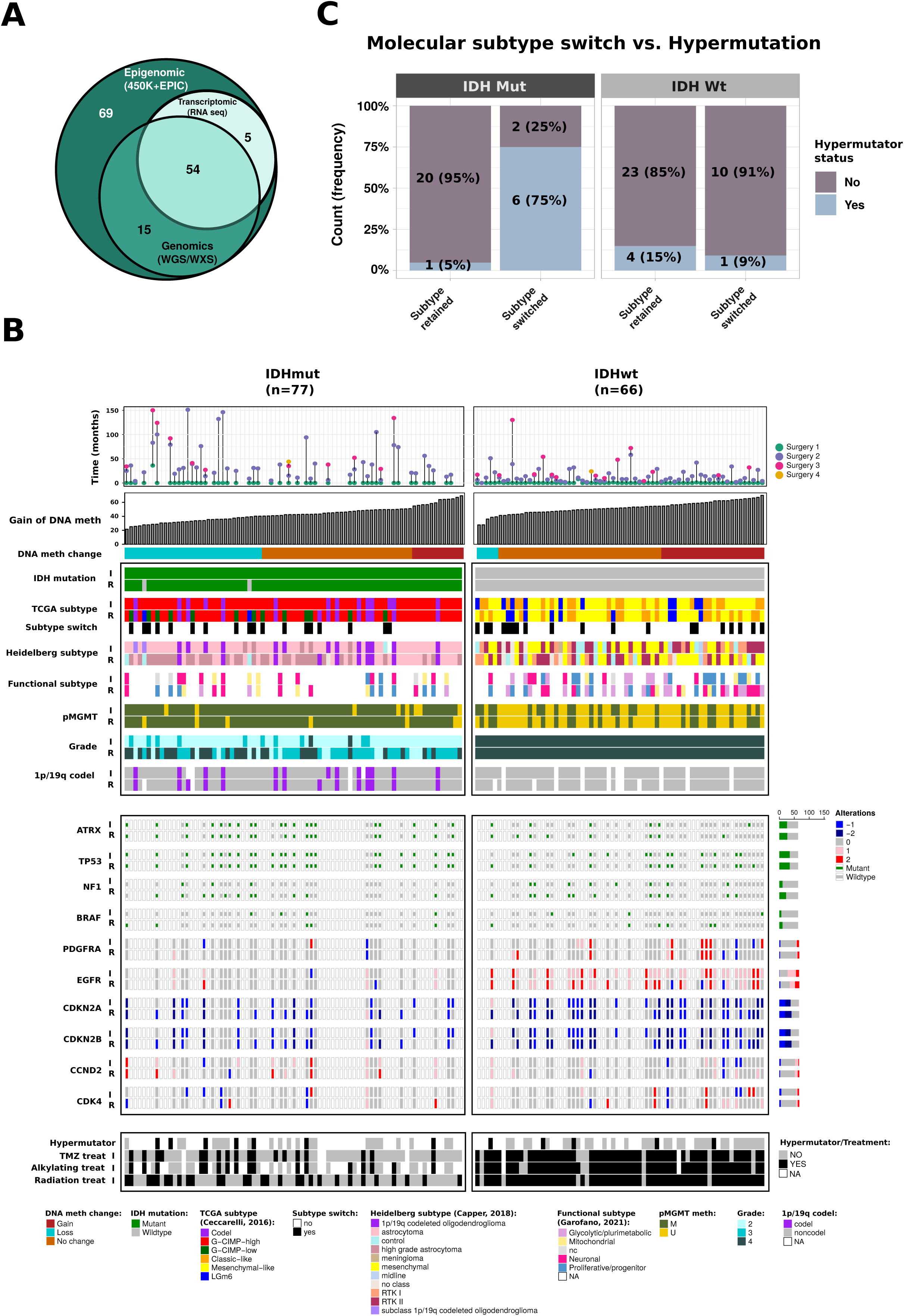
Epigenomic evolution of matched initial and recurrent gliomas. **A)** Venn diagram of number of patients who had DNA methylation, genomic (WGS/WXS) and/or RNAseq profiling. **B)** Clinical and molecular overview of matched initial and recurrent DNA methylation cohort. Each column represents a single patient (n=143) at two separate time points grouped by IDH status and ordered by increase of gain of DNA methylation at recurrent tumor from left-to-right. Top plot shows the surgical interval of each patient. **C)** Frequency of patients with hypermutator tumors that switched or retained molecular subtype at recurrence. Patients are divided by IDH status.

To investigate the temporal differences we evaluated the most relevant molecular and clinical features in gliomas. Our cohort includes the previously described DNA methylation-based glioma TCGA subtypes: Three IDH-mut-specific DNA methylation subtypes (Codel, G-CIMP-high, and G-CIMP-low) and three IDHwt-specific subtypes (Classic-like, Mesenchymal-like, LGm6) (Ceccarelli et al., 2016) (Figure 1B) and we used this molecular classification throughout the analyses in this study. Over time, 25% (19/77) of IDHmut and 30% (20/66) of IDHwt tumors switched subtypes. Among IDHmut cases that switched subtypes, tumors of 13 patients (13/19, 68%) switched from G-CIMP-high to G-CIMP-low, a subtype associated with worse overall survival (time from initial surgery to death or last follow-up). Among IDHwt cases that switched subtype, tumors of 10 patients (10/20, 50%) switched from classic-like or LGm6 to the mesenchymal-like subtype. Less frequent subtype shifts were also observed. A Pan-CNS DNA methylation-based classification (Heidelberg subtype; Capper et al., 2018) and a recent pathway-based classification of glioblastomas (Functional subtype; Garofano et al., 2021) were also assigned to our cohort (Figure 1B). Based on the Pan-CNS DNA methylation-based classification, 26 IDHmut astrocytomas (26/55, 47%) progressed to high-grade astrocytoma upon recurrence; whereas the IDHwt cases, tumors of 7 patients (7/22, 32%) switched from mesenchymal to RTK II, 6 cases (6/22, 27%) switched from RTK II to mesenchymal and 3 (3/9, 33%) from RTK I to mesenchymal.

When the patients were stratified according to the genome-wide gain or loss of DNA methylation groups upon recurrence, patients with IDHmut gliomas showed a higher proportion of samples losing DNA methylation than patients with IDHwt gliomas (42% vs 6%; Fisher’s test, p < 0.0001; Figure 1B). Interestingly, 6 of 8 (75%) of IDHmut tumors that switched molecular subtypes and had genomic data available evidenced hypermutator phenotype at recurrence (Figure 1C, Table S3B). Five of these switched from G-CIMP-high to G-CIMP-low. The tumor of one patient switched from IDHmut Codel to G-CIMP-high (non-codel subtype). In contrast, only 1 of 21 (5%) IDHmut patients which retained their subtype became hypermutator at recurrence (Fisher’s test, p=0.0004). All switches were toward a more aggressive phenotype (e.g. G-CIMP-low and grade 4), suggesting an association between DNA methylation change, tumor progression, and hypermutation acquisition.

To further explore the association of epigenomic changes and hypermutation, we investigated the methylome of the initial tumors of patients that subsequently received chemotherapy, particularly TMZ, to identify biomarkers that could predict hypermutation in gliomas. We identified 6 IDHmut and 4 IDHwt initial tumors that transformed to hypermutators upon the first recurrence and had treatment information available (Figure S1A). For comparison, we identified 5 IDHmut and 5 IDHwt samples that, despite receiving TMZ and/or RT after initial surgery, did not become hypermutators which were used as controls.

A supervised analysis between initial IDHmut gliomas that did or did not develop a hypermutator phenotype at recurrence resulted in 342 differentially methylated CpG probes (unadjusted p-value < 0.01 and DNA methylation difference > 25%) (Figure S1B, Table S5). To understand the biological implications of these DNA methylation changes, we integrated the methylome data with RNA-sequencing and selected CpGs within the promoter region of genes and identified 23 CpG-gene pairs inversely correlated, such as the DNA replication initiator gene *REPIN1* and the gene *KCNK2,* which has been associated with positive regulation of cellular response to hypoxia (Wu et al. 2013), indicating potential mechanisms associated with the predisposition to hypermutation (Figure S1C). Within IDHwt tumors, we identified 548 CpG probes that were associated with hypermutation in IDHwt gliomas (unadjusted p-value < 0.05 and DNA methylation difference > 30%) and 14 genes being negatively correlated with CpG methylation, including *MGMT* and the gene *NETO1*, which is associated with regulation of long-term neuronal synaptic plasticity (Ng et al. 2009) (Figure S1D and S1E, Table S5). Further studies would be needed to confirm the predictive power of these signatures.

Despite evidence that hypermutation might occur in random chromosomal locations (Touat et al., 2020), we investigated the methylome of the recurrent hypermutated tumors to identify regions with consistent patterns of DNA methylation. We compared the epigenome between first recurrence hypermutators (N=8 IDHmut; 7 IDHwt) and non-hypermutators (N=11 IDHmut; 20 IDHwt), stratified by IDH status. In the IDHmut group, we identified 502 differentially methylated CpG probes (DMP) (unadjusted p-value < 0.01 and DNA methylation difference > 30%) that defined the hypermutator phenotype in gliomas, the majority of which were hypomethylated (Figure S1F, Table S5). In IDHwt tumors, the set of CpG probes identified by our analysis (N=620; unadjusted p-value < 0.05 and DNA methylation difference > 40%) showed that the hypermutators are mostly hypermethylated compared to non-hypermutators (N=94%; 581/620) (Figure S1G, Table S5). Taken together, these findings suggest that epigenetic biomarkers predict the predisposition to hypermutation in patients who receive TMZ after the initial surgery. Further validation studies could help elucidate the biological role of these regions in determining the susceptibility to hypermutation.

### Master regulators associated with IDHmut glioma progression

To further investigate the changes in the epigenome that occur over time in gliomas, we compared the genome-wide DNA methylation characteristics of the initial compared to recurrent tumor samples, stratified by IDH status (Figure S2B). IDHwt gliomas showed a more stable epigenome over time (i.e. zero CpG probes presented a differentially methylated mean difference greater than 20%), while the epigenome of IDHmut gliomas showed genome-wide loss of DNA methylation (593 CpG probes with DNA methylation difference > 20%) throughout the disease evolution (Figure S2B). IDHmut patients that progressed from G-CIMP-high to G-CIMP-low showed the most prominent loss of DNA methylation, particularly at intergenic regions, which suggests an association between the loss of DNA methylation and tumor progression in this subtype (Figures S2C, S2D and S2E), confirming previous results from our group and others (Nomura et al., 2019; de Souza et al., 2018). Compared to patients with tumors that remained G-CIMP-high at recurrence, those with recurrent G-CIMP-low more often had histologically higher-grade astrocytoma, were less often managed by a watch-and-wait strategy and exhibited inferior survival (Table S3A).

Next, we investigated the impact of the epigenomic changes on transcriptional networks using an integrative approach that combines epigenome and transcriptome to define master regulators. We applied ELMER (Enhancer Linking by Methylation/Expression Relationships) (Silva et al., 2019; Yao et al., 2015), a computational tool that harnesses DNA methylation to identify regulatory elements and correlates the enhancer state with the expression of nearby genes to define putative transcriptional targets. ELMER also allows the inference of transcription factor (TF) binding motif analysis and the integration of TF expression to identify master regulatory TFs (MRTFs) (Silva et al., 2019).

We compared initial and recurrent IDHmut gliomas and identified a total of 1,590 DMP between the groups (FDR < 0.05, Figure 2A, S4A). To better understand the genomic context of the DNA methylation changes associated with the progression of IDHmut gliomas, we profiled nine IDHmut G-CIMP-high gliomas at diagnosis and three recurrences evolving into IDHmut G-CIMP-low states with ChIP-seq for the H3K27Ac active chromatin mark. The integration of DMP with the local changes of H3K27Ac (20 nearest genes of each CpG probe in the methylation array) converged on the identification of 1,282 genes exhibiting coordinated loss of DNA methylation and gain of H3K27Ac, thus indicating that this gene set is epigenetically activated by loss of DNA methylation at recurrence. A pathway-based analysis of these genes suggested enrichment of cell cycle and proliferation-related activities (Figure 2B, Table S4B).

**Figure 2.**
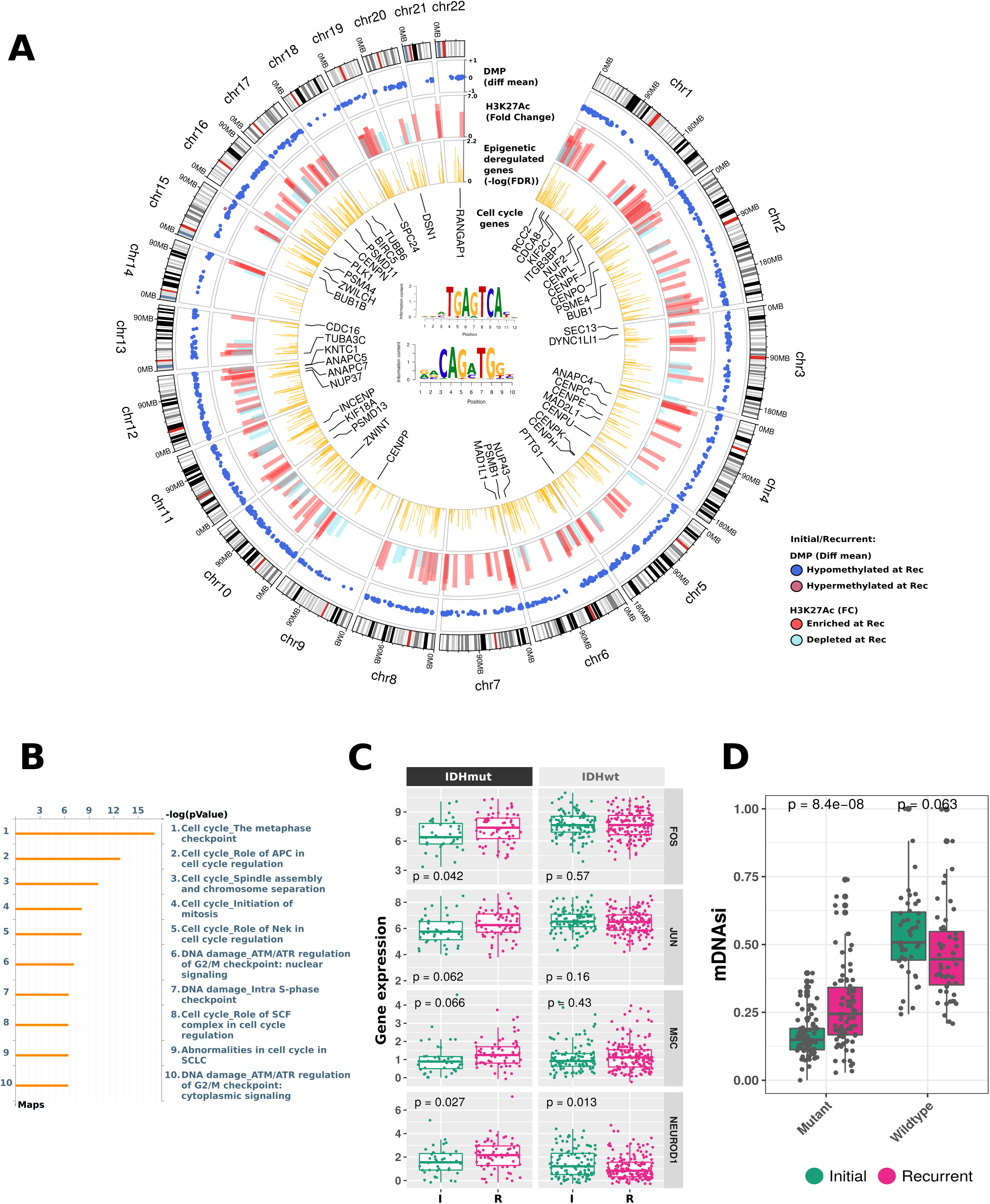
Master regulators associated with IDHmut glioma recurrence/progression. **A)** Circos plot diagram showing epigenomic changes between initial and recurrent IDHmut tumors. The outer track shows chromosome location. The second track shows genomic location of 1,590 differentially methylated CpG probes between initial and recurrent IDHmut tumors. Height of dots represents the difference of methylation levels between recurrent and initial. The third track shows the genomic location of H3H27ac peaks defined by differential binding analysis of IDHmut G-CIMP-high patients that progressed to G-CIMP-low. Height of the peaks represents the fold change of differential binding analysis. The fourth inner track shows the genomic location of the candidate genes epigenetically regulated by the CpG probes (yellow bars). Height of the bars represents the -log of FDR. Genes related to cell cycle are labeled. The top transcription factor (TF) binding motifs enriched among the candidate DMP are FOS/JUN family and NEUROD1, whose DNA signature motifs are represented in the center of the circos plot. **B)** Pathway analysis of epigenetically activated genes in recurrent IDHmut tumors. **C)** Gene expression levels of master regulator transcription factors (MRTFs) of IDHmut initial and recurrent tumors. IDHwt tumors are shown as a reference. **D)** Stemness activity in glioma samples stratified by primary/recurrent and IDH mutation status.

Within the regulatory DMP regions, we identified the enrichment of 32 distal-binding motifs for specific TFs with TGAGTCA (Fos/Jun family) and the enhancer-box CAGATGG being the most represented (Figure 2A, Table S4C). By computing the anticorrelation of TF gene expression and the DNA methylation levels within their binding sites, we identified candidate TFs with increasing expression in IDHmut recurrent tumors to levels comparable to those of IDHwt gliomas (Figure 2C). Some of these master regulatory (MR) TFs are well-established glioma oncogenes: FOS/JUN family members have been associated with control of tumor malignancy (Blau et al., 2012), cell proliferation and radioresistance (Liu et al., 2016), and glioma progression (de Souza et al., 2018); musculin (MSC) has been reported to inhibit differentiation of embryonal carcinoma (Yu et al., 2003) and B-cell lymphoma cells (Mathas et al., 2006). Indeed, recurrent IDHmut tumors show higher stemness activity (as defined by our recent study) (Malta et al., 2018b) than the corresponding initial tumors (Figure 2D).

We also identified 201 DMP between initial and recurrent IDHwt glioma samples, which were mapped to both distal regions (156 CpG probes) and promoters (45 CpG probes) and their putative associated genes (Figure S2F and S2G). We did not identify any TF binding motifs enriched within the DMP regions, showing again less frequent changes in epigenetic regulation in IDHwt tumors over time.

Together, our results show that IDHmut gliomas present a more dynamic epigenome, characterized by loss of DNA methylation at recurrence, with activation of oncogenes to expression levels similar to those observed in IDHwt gliomas. In contrast, the epigenome of IDHwt gliomas seems relatively preserved longitudinally. We identified potential master regulators driving cell cycle deregulation in IDHmut gliomas at recurrence, which represent potential treatment targets.

### DNA methylation loss associated with recurrent IDHmut gliomas after standard treatment

It has been shown that treatment (radiotherapy and alkylating chemotherapy) improves the progression-free survival of IDHmut gliomas (Wick et al., 2009). However, recurrence of IDHmut lower-grade glioma is frequently associated with progression to higher histological grades. Treatment of IDHmut lower-grade gliomas with TMZ and/or radiotherapy has been linked to many genomic alterations, such as a hypermutator phenotype (Johnson et al., 2014; Touat et al., 2020), and a strong tendency towards aneuploidy and a specific radiotherapy-associated deletion signature by genetic analysis (Kocakavuk et al., 2021). Herein, we observe that treatment is associated with epigenomic changes. Whether and how to treat a low-grade IDH-mutant glioma is still not based on high level evidence from controlled clinical trials; we sought to identify the DNA methylation changes triggered at recurrence by the different treatment choices made in our cohort. Towards this goal, we divided our cohort into 4 groups: patients who received TMZ only (N=12), radiotherapy (RT) only (N=18), the combination of RT and TMZ (RT+TMZ; N=6) and patients who did not receive additional treatment after the first surgery but were managed by a watch-and-wait approach (N=33) (Tables S3D and S6). The supervised methylome analysis of first recurrent IDHmut gliomas across the 4 groups defined 620 DMP (Kruskal-Wallis test by ranks, FDR < 0.01 and DNA methylation difference > 20%) (Figure 3A, Table S6). Upon first investigation, we determined that these CpGs were associated with consistent loss of DNA methylation in patients who received any treatment besides surgery after initial diagnosis (TMZ only, RT only or combined TMZ and RT) compared to their primary counterparts. On the other hand, the methylome of the recurrent sample of patients who did not receive treatment resembled the initial tumor (Figure 3A). There were no CpG probes that distinguished the different groups that received any specific type of treatment (TMZ only, RT only or RT+TMZ), suggesting that the introduction of any treatment regimen involving TMZ and/or RT after initial surgery is sufficient to remodel the methylome of treatment-*naïve* gliomas. The interrogation of the same CpGs corroborated this observation in IDHwt initial and recurrent gliomas, which showed a similar DNA hypomethylation profile to the IDHmut treatment arm (TMZ only, RT only and RT+TMZ) (Figure 3A). To evaluate whether the observed decrease of DNA methylation in our discovery cohort (GLASS-International; this study) is consistent in an independent cohort, we sought to validate our findings in a yet unpublished dataset of paired glioma samples from the GLASS in the Netherlands consortium (GLASS-NL; validation) (GLASS Consortium, 2018). The validation cohort consists of 36 treated paired glioma samples and 64 untreated paired samples, all of which are IDHmut astrocytomas at diagnosis. The loss of DNA methylation pattern upon treatment after initial surgery was confirmed in the validation cohort (Figures 3B and 3C). Because the GLASS-NL cohort was comprised of only IDHmut astrocytomas, we repeated our discovery test on only the IDHmut astrocytomas of the GLASS-International cohort (excluding oligodendroglioma samples), separated by treatment group and selected a new set of DNA hypomethylated CpG probes (N=981) in the treated samples (FDR < 0.01 and DNA methylation difference > 25%). 61% (381/620) of the previously described CpG probe list overlapped with this new signature showing consistent DNA methylation changes. Again, the loss of DNA methylation pattern after treatment was confirmed in the validation cohort (Figure S3A, Table S6).

**Figure 3.**
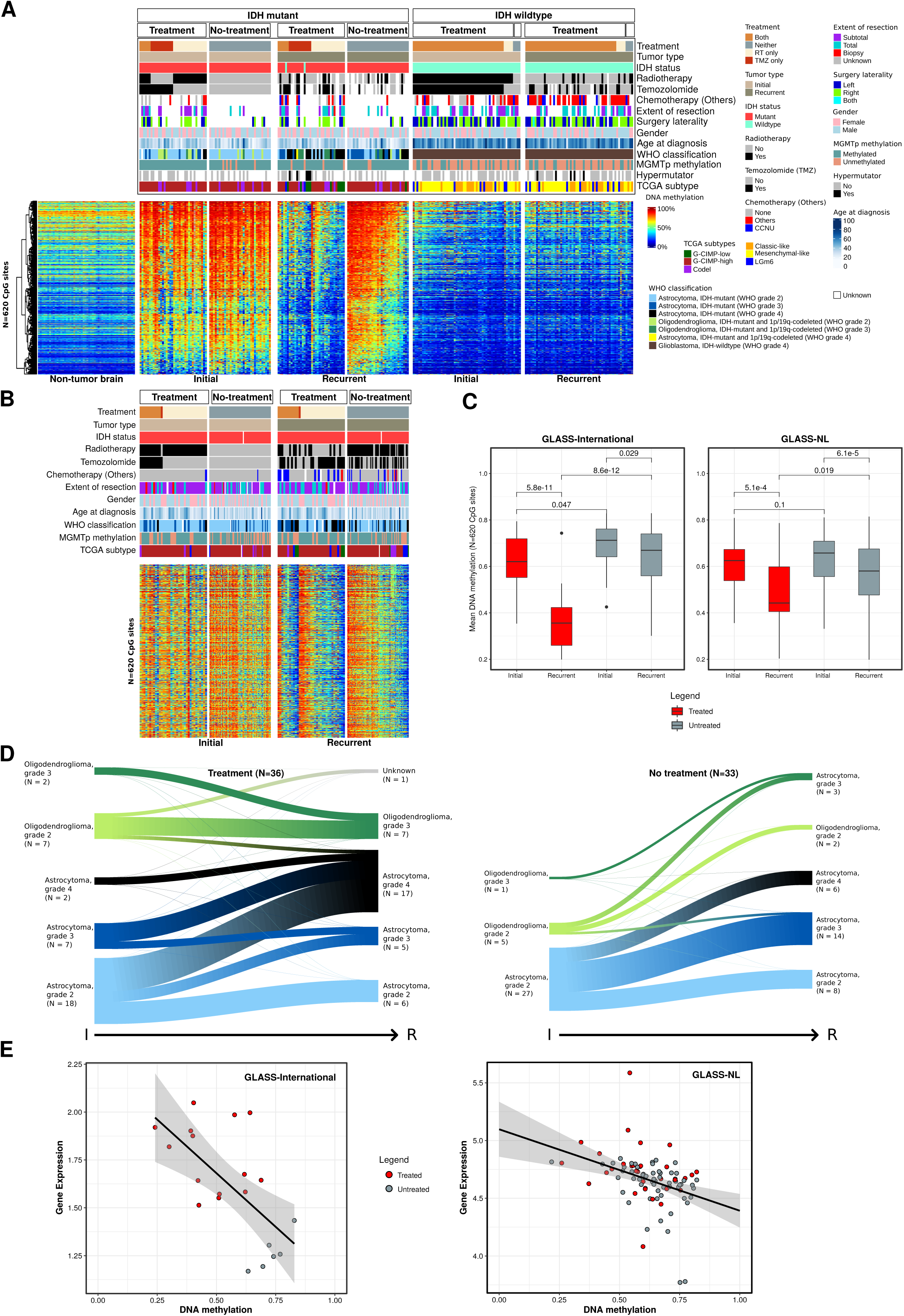
DNA methylation loss associates with malignant transformation of glioma after standard treatment. **A)** Heatmap of DNA methylation data. Supervised hierarchical clustering analysis of 620 CpG probes that are associated with different treatment strategies in IDH-mutant paired glioma samples. Columns represent glioma samples, rows represent CpG probes. Samples were stratified and clustered based on IDH mutation status and primary/recurrent status and CpGs were ordered using hierarchical clustering methods. Non-neoplastic brain samples are represented on the left of the heatmap. DNA methylation beta-values range from 0 (low) to 1 (high). Additional tracks are included at the top of the heatmaps to identify each sample membership within separate cluster analysis. **B)** Heatmap of DNA methylation data in the validation cohort - GLASS-NL, showing the same 620 CpG probes of panel A. **C)** Boxplot of the average DNA methylation beta-value of the 620 CpG probes from panel A. Samples are stratified by primary/recurrent status and by treated/non-treated status. Left: GLASS-International samples. Right: GLASS-NL samples. **D)** Evolution of tumor histology (2021 WHO classification) from primary to recurrent samples after treatment compared to non-treated gliomas. **E)** Scatter plot of mean DNA methylation of CpG probes and mean gene expression of the epigenetically regulated genes after treatment.

By comparing different clinical and molecular features of these tumors, we observed a significant enrichment of progression to IDHmut astrocytoma CNS WHO grade 4 in the treatment vs the non-treatment groups in the discovery (Fisher’s test = 0.03; Figure 3D) but not with the validation datasets (Fisher’s test p = 0.3; Figure S3B). In addition, we noticed that 34% (10/29) of G-CIMP-high tumors progressed to G-CIMP-low in the treatment group vs 4% (1/27) in the non-treatment group (Fisher’s test, p=0.005). The same was observed with the molecular Pan-CNS classification (Capper et al., 2018), in which 68% (15/22) of IDHmut astrocytomas progressed to high-grade astrocytoma in the treatment group vs 23% (6/26) in the non-treatment group (Fisher’s test, p=0.003, Table S1). As expected, only in the treatment group, we observed hypermutator samples (N=5) (Figure 3A). To consider whether the selection of the treatment regimen might have been biased by the age at diagnosis, we performed an analysis of variance (ANOVA) on age at the initial tumor resection and we did not find an age difference across the four groups (ANOVA F-test p-value = 0.59). To explore the biological implications of these epigenomic changes, we searched for inversely correlated DNA methylation and gene expression. We identified 24 distal CpG-gene pairs (FDR < 0.01) enriched for the NEUROD1 motif resulting in 18 unique genes potentially regulated by DNA methylation after treatment (Figures 3E and S3C, Table S6) which included known oncogenes and cancer-related genes such as MYB and MYO15A and enriched for monocarboxylic acid metabolic process.

### Tumor microenvironment changes and clinical implications of treatment in IDH-mutant gliomas revealed by DNA methylation

As the observed changes of DNA methylation may not only be driven by tumor cell-intrinsic-events but may also underscore changes in the methylome associated with non-tumor cell populations present in the glioma tumor microenvironment (TME), we applied a methylation-specific approach for the deconvolution of non-tumor cells to the longitudinal glioma cohort. MethylCIBERSORT (Chakravarthy et al., 2018) uses genome-wide DNA methylation data to deconvolute cell types from individual tumor samples. Our estimated cell populations consisted of 10 cell types: glia, neuron, endothelial cells (CD31+), B cells (CD19+), CD4 effector T cells, CD8+ T cells, T regs, natural killer cells (CD56+), macrophages and neutrophils. Compared to untreated recurrencies, recurrent IDHmut tumors which were treated after surgery were marked by increased infiltration of endothelial cells (CD31+) and CD8 T lymphocytes (CD8+) (Figure 4A, Table S7), suggesting that treatment impacts the tumor microenvironment and most notably angiogenesis. Our *in silico* cell fraction estimation was validated by immunohistochemical staining in representative IDHmut samples that received treatment after initial surgery (Figure 4B). As a further validation, the relative proportion of non-tumor cells estimated by methylCIBERSORT was significantly correlated with the glioma cell compartments estimated by gene expression for the GLASS-international transcriptomic cohort (Varn et al., 2021) (Figure S4A). Additional changes in the TME were also observed within IDHmut and IDHwt longitudinally, particularly a significant increase in macrophage in recurrent IDHwt tumors (Figures S4B-S4D).

**Figure 4.**
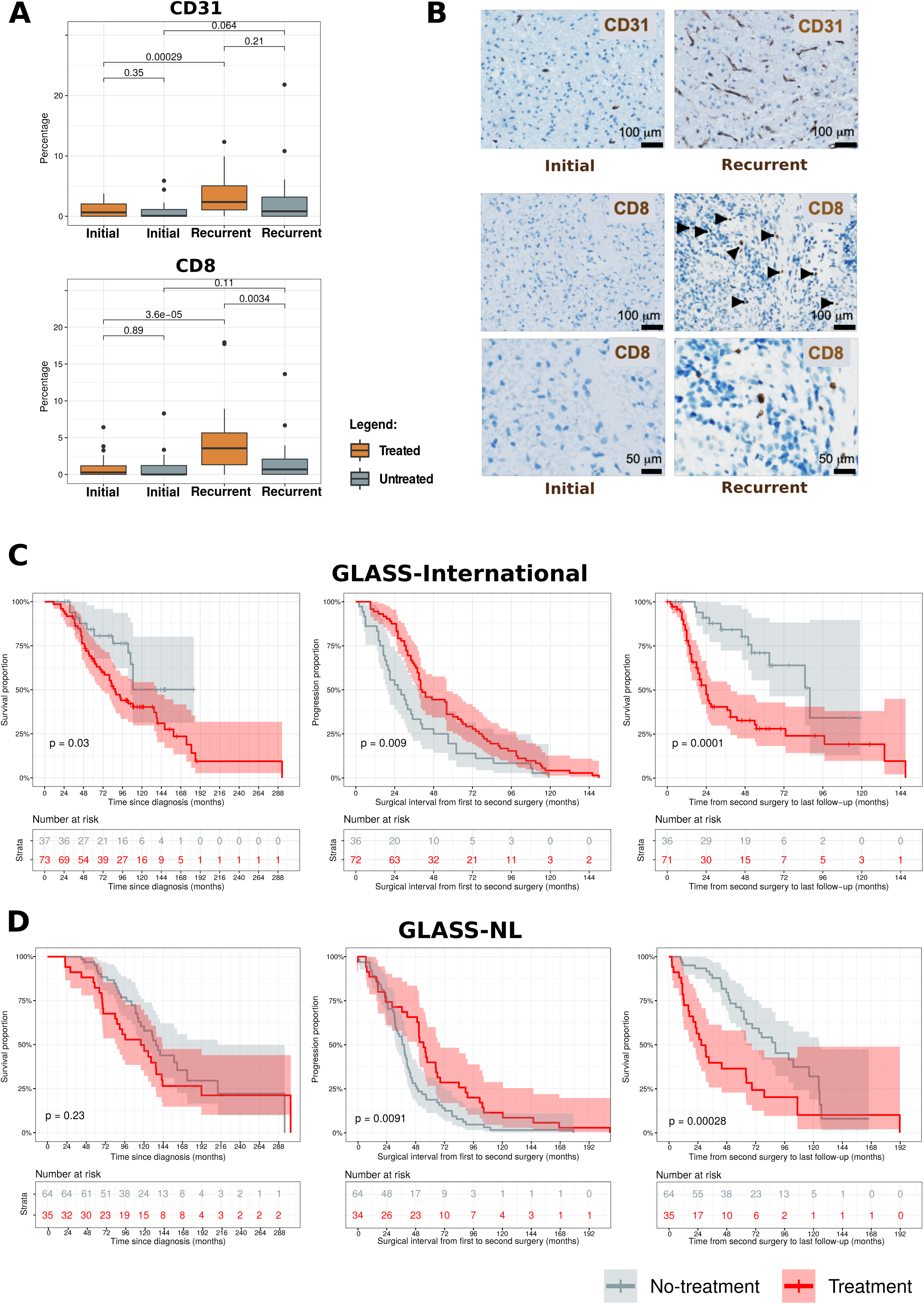
Tumor microenvironment and clinical implications of treatment in IDH-mutant gliomas. **A)** CD31 and CD8 proportions (range scaled from 0 to 100%) in samples originating from IDHmut matched initial and recurrent tumors in treated and non-treated patients. **B)** Illustrative immunohistochemical stainings for two marker proteins (CD31 and CD8) in an individual patient showing change of levels of tumor-infiltrating immune cells between initial and recurrent tumors. **C/D)** Overall survival and surgical interval analysis of IDHmut gliomas for the GLASS International (C) and GLASS-NL (D) cohorts.

To evaluate the clinical implications of these findings, we next assessed the clinical follow-up of the entire IDHmut GLASS-International cohort divided into treatment (N=73) or non-treatment (N=37) groups. First, patients receiving treatment after initial surgery had a worse survival than patients who did not receive treatment beyond surgery (log-rank p-value=0.03, Figure 4C, left). However, patients left untreated after initial surgery had a progression-free interval (PFI) of 27 months that was significantly worse compared to a PFI of 40.5 months in the treatment group (log-rank p=0.009, Figure 4C, middle), thus confirming the results of EORTC 22845 (van den Bent et al., 2005). When we focused on the survival interval from second surgery to the last follow-up, as expected, we found a markedly worse survival in previously treated patients (log-rank p=0.0001, Figure 4C, right). The same associations of treatment and clinical implications were observed in the validation GLASS-NL cohort (Figure 4D).

## Discussion

This work reports and analyzes the largest cohort of matched glioma samples profiled with epigenomics, transcriptomics, and genomics platforms to uncover the diverse molecular routes that drive treatment-resistant gliomas during progression. By applying an integrative molecular approach, we highlight the critical epigenetic mechanisms by which gliomas evaded treatment in addition to transcriptomic and genetic evolution. We also identified epigenomic changes that may be useful as biomarkers for continuous monitoring of disease progression and treatment response prediction.

Our cohort consisted of the three major glioma subtypes (IDHmut-noncodel, IDHmut-codel and IDHwt) and revealed key evolutionary differences across these subgroups. Earlier work from us and others described the G-CIMP glioma phenotype in IDHmut glioma characterized by higher levels of DNA methylation likely as a direct result of the epigenetic effects of the oncometabolite 2-hydroxyglutarate that is generated by mutant IDH enzymes (Noushmehr et al., 2010; Turcan et al., 2012). These tumors exhibited favorable clinical outcomes compared with IDHwt gliomas, which have lower DNA methylation levels (Noushmehr et al., 2010). Indeed, in our previous work, we reported that the extent of genome-wide DNA methylation showed broad positive correlation with clinical outcome for IDHwt glioma, now defined as “molecular GBM’’ given their aggressive behavior independent of histological grading (Louis et al., 2001), with these tumors having the lowest levels of DNA methylation genome-wide. Here we confirm and extend these notions to tumor progression. In particular, we found that IDHwt gliomas presented an initial low genome-wide DNA methylation, which shows minimal changes during the course of the disease. This is consistent with the malignant state of IDHwt gliomas at diagnosis and which does not appear to progress during treatment.

Conversely, we uncovered pronounced epigenetic changes in IDHmut gliomas, which invariably converged toward lower DNA methylation levels in recurrent treated tumors compared to untreated neoplasms. In the most extreme cases, the treatment-induced evolution of IDHmut glioma resulted in a state of DNA hypomethylation comparable to IDHwt gliomas. Thus, the epigenetic trajectory of glioma progression was associated with progressively lower levels of DNA methylation in IDHmut tumors or was essentially moot in the case of IDHwt gliomas as glioma initiation in the absence of IDH mutations coincided with the lowest possible levels of DNA methylation.

Following diagnosis and surgery, IDHmut glioma patients may or may not undergo treatment with radiation or alkylating chemotherapy or both (Baumert et al., 2016; Buckner et al., 2016). A recognized effect of alkylating agents typically used in glioma therapy (temozolomide) is the potential acquisition of a hypermutator phenotype (Barthel et al., 2019; Johnson et al., 2014). Here, we found that treatment with radiation and/or chemotherapy individually or combined accelerated progression towards the hypomethylated state at recurrence. These findings were confirmed in an independent cohort, suggesting that current treatment regimens alter the epigenomic landscape of IDHmut gliomas similar to the observed treatment-induced genomic abnormalities in response to alkylating agents.

The treatment-accelerated epigenetic drift towards the hypomethylated state parallels the histopathological shift from a lower-to a higher-grade phenotype. Altogether, these results indicate that the initial treatment with radiotherapy and/or temozolomide may have triggered a more aggressive evolution at the time of the tumor recurrence which compromised the survival probability of these patients at that recurrence. More specifically, the introduction of treatment after initial surgery in IDH-mutant gliomas is associated with a significant delay in tumor progression. The time from second surgery to progression or death is significantly shorter when compared to untreated patients which is likely associated with the loss of DNA methylation and activation of associated genes that we uncovered. While these findings remain consistent with previous observations from large clinical trials that reported the beneficial role of chemotherapy and radiotherapy for patients survival (van den Bent et al., 2005), they also highlight the ability of radio/chemotherapy to trigger a markedly more aggressive behavior at recurrence and identify the epigenetic determinants of the treatment-induced glioma progression.

The therapy-associated changes of the epigenetic evolution of IDHmut glioma were also mirrored by specific changes in the tumor microenvironment (TME) at recurrence. At recurrence, CD8 and endothelial cell-related signatures were elevated in treated IDHmut gliomas compared to untreated tumors. Together, these findings indicate that the epigenomic and genomic changes associated with more aggressive histotypes of IDHmut gliomas at recurrence coincided with specific changes of the TME (neo-angiogenesis and changes in T-cell composition), indicating the convergence of IDHmut glioma evolution towards features more typical to the most aggressive IDHwt subtype.

Having established the importance of epigenetic evolution of IDHmut glioma, we focused on this subtype to integrate the analysis of epigenome, transcriptome and ChIP-seq to extract the features associated with the evolution towards loss of DNA methylation and consequent transcriptional activation of critical drivers of progression (master regulators, MRs). First, the integration of DNA methylation and transcriptomic platforms uncovered the convergence towards gene sets enriched with cell cycle and proliferation-related activities and stemness, in comparison to initial tumors. Thus, such biological activities are directly activated by transcriptional derepression associated with local loss of DNA methylation. Second, we focused our attention on identifying MRs that enabled the biological evolution of IDHmut glioma. MR are TFs regulating multiple genes and in the case of cancer, drive the onset or progression of the disease.

Our approach depends on our understanding that discrete regions of the genome harbors DNA sequence elements recognized by TFs. The presence of the TF and the accessibility of the functional elements can then drive the activity of nearby or distant genes. We uncovered a set of established oncogenes as MRs activated by hypomethylation-induced derepression in IDHmut glioma, among them FOS/JUN family members. Interestingly, therapeutic targeting of cJun transcription has included small molecules interfering with its DNA binding domain, thereby preventing it from activating downstream target genes such as those represented in our study. Two such molecules with potential therapeutic advantages are MLN944 (Dai et al., 2004) and retinoid (SR11302) which could be explored in a follow-up study involving recurrent glioma tumors.

In summary, we found that upon tumor progression after standard treatment loss of DNA methylation in IDHmut tumors is frequent which results in epigenetic activation of cell cycle-related genes associated with early tumor progression and alterations in the TME contexture towards angiogenesis and a T-cell composition that resembles a treatment-naive IDHwt glioma (Figure 5). In untreated IDHmut patients the epigenome does not change significantly, and tumors progress later than the treated ones, either spontaneously or after subsequent treatment.

**Figure 5.**
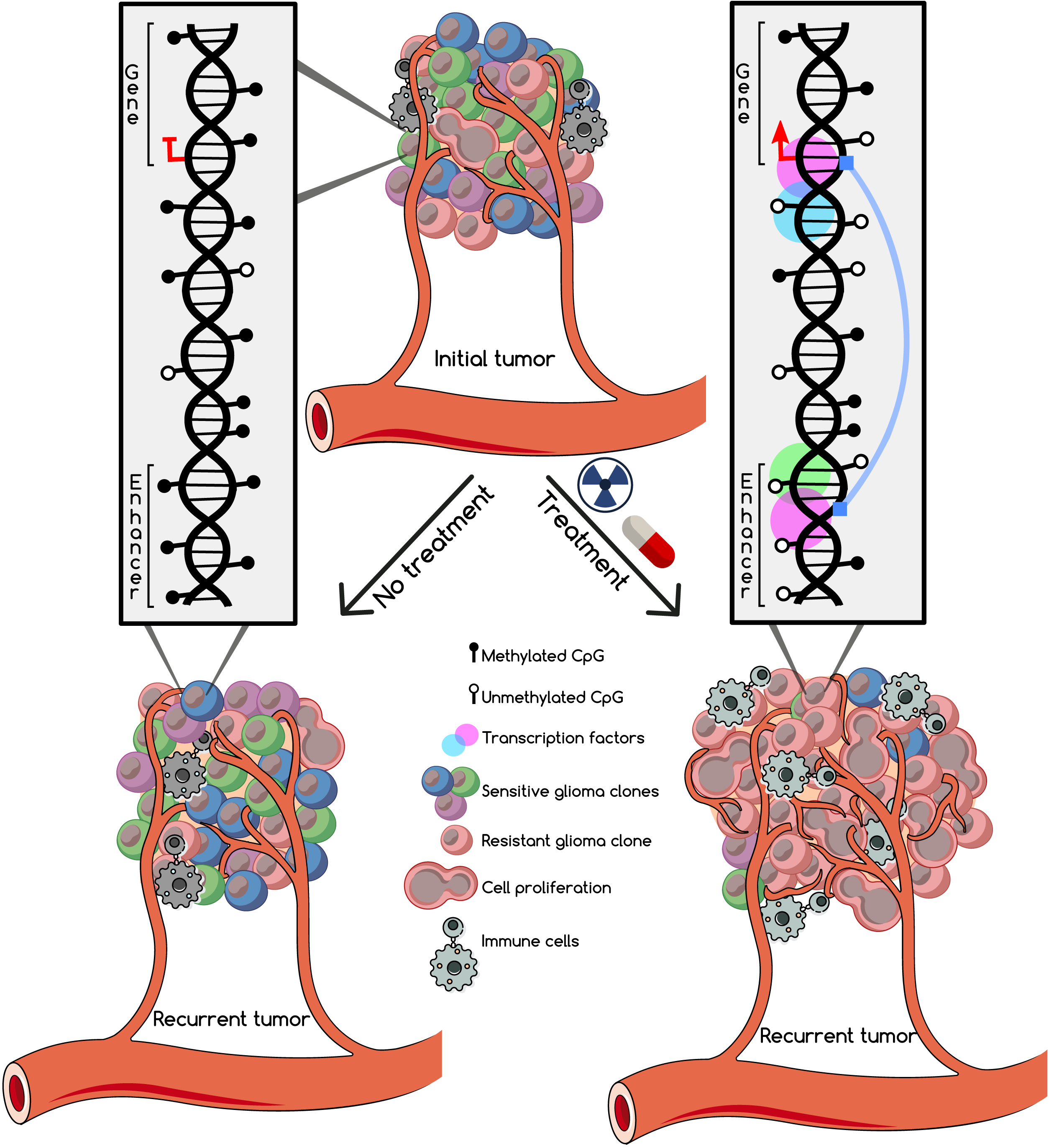
Schematic of treatment-induced epigenetic activation of cell cycle in IDHmut recurrent tumors. Standard treatment induces loss of methylation in IDHmut tumors with further epigenetic activation of cell cycle related genes associated with early tumor progression. In IDHmut non-treated patients, epigenetic changes are much more limited and tumors may progress later, either spontaneously or after subsequent treatment.

## Supplemental Figure Legends

**Figure S1: DNA methylation signature to predict the hypermutator phenotype upon initial surgery divided by IDH status, related to Figure 1. A)** Venn diagram of the GLASS hypermethylator cohort based on data type: Whole-genome sequencing, DNA methylation array and clinical follow-up. **B)** Heatmap of DNA methylation data across IDHmut samples at initial diagnosis that were treated with TMZ, stratified by hypermutator status. Supervised hierarchical clustering analysis of 342 CpG probes that can predict the hypermutator status at the initial surgery. Column-wise represents 11 glioma samples, row-wise represents CpG probes. Samples and CpGs are ordered using hierarchical clustering method. DNA methylation beta-values range from 0 (low) to 1 (high). Additional tracks are included at the top of the heatmaps to identify each sample membership within separate cluster analysis. **C)** Boxplot of the CpG-gene pairs at gene promoter regions of IDHmut gliomas divided by DNA methylation (top) and gene expression (bottom) data. **D)** DNA methylation heatmap of 548 CpG data across IDHwt samples at initial diagnosis treated with TMZ, stratified by hypermutator status. **E)** Boxplot of CpG-gene pairs at gene promoter regions identified in IDHwt gliomas. **F/G)** Differentially methylated regions between hypermutator vs non-hypermutator recurrent samples divided by IDH status: F, IDHmut and G, IDHwt.

**Figure S2. Longitudinal epigenomic changes of matched initial and recurrent gliomas, related to Figure 2. A)** Venn diagram of DNA methylation samples which were profiled with genomics (WGS/WXS) and/or RNAseq. **B)** Overall genome-wide DNA methylation correlation between initial and recurrent tumors, stratified by IDH mutation status. **C/D/E)** Overall DNA methylation correlation between initial and recurrent tumors of different glioma subtypes, by CpG probe genomic location: C, genome-wide; D, probes located in intergenic regions, and E, CpG probes located in promoters of genes. **F/G)** Differentially methylated CpG probes defined between recurrent and initial IDHwt tumors and their associated putative genes. F, CpG probes located at distal regions and G, CpG probes located at promoter regions.

**Figure S3. DNA methylation loss is associated with malignant progression of glioma after standard treatment in the validation cohort: GLASS-NL, related to Figure 3. A)** Heatmap of DNA methylation data. Supervised analysis using astrocytoma-only samples from GLASS-International cohort identified 981 CpG probes that are associated with treatment astrocytomas IDHmut paired glioma samples. Samples from the validation cohort are also shown. Samples are stratified by cohort, initial/recurrent status and treatment status. Column-wise represents glioma samples, row-wise represents CpG probes. DNA methylation beta-values range from 0 (low) to 1 (high). Additional tracks are included at the top of the heatmaps to identify each sample membership within separate cluster analysis. **B)** Evolution of tumor histology (2021 WHO classification) of the validation cohort (GLASS-NL) after treatment compared to non-treated gliomas. **C)** Scatter plot of 24 CpG-gene pairs epigenetically regulated genes after treatment in IDHmut gliomas.

**Figure S4. Glioma subtypes present different tumor microenvironments and it changes overtime, related to Figure 4. A)** Correlation between cell composition estimated by DNA methylation and by gene expression. **B)** Barplots of the estimated median infiltration of specific cell types as a proportion of all cell types (range scaled from 0 to 100%) in 143 glioma tumors at initial diagnosis, divided by IDH mutation status. **C)** Cell type proportion (range scaled from 0 to 100%) in samples originating from the matched initial and recurrent tumors, divided by molecular subtypes. All comparisons (initial vs. recurrence, by subtype, for the specified cell population) are statistically significant (P < 0.05). P values calculated using a two-sided Wilcoxon rank-sum test. Matched primary and recurrent tumors are linked by the lines. **D)** Representative immunohistochemical staining for CD163 marker protein in two individual patients with changing levels of tumor-infiltrating immune cells between initial and recurrent tumors.

## Methods

### CONTACT FOR REAGENT AND RESOURCE SHARING

Further information and requests for resources and reagents should be directed to and will be fulfilled by the Lead Contact Houtan Noushmehr.

#### Biospecimens/GLASS Datasets

Datasets added to GLASS came from both published and unpublished sources. The GLASS epigenomic cohort consists of 357 DNA methylation samples (143 patients) profiled by either Illumina 450K or EPIC Beadchip methylation arrays and described below. For those same patients, we also collected DNA sequencing data of 157 genomic samples, WXS or WGS (69 patients); and RNA sequencing of 120 samples (59 patients), available through the GLASS consortium, resulting in the largest cohort of matched glioma samples profiled with epigenomics, transcriptomics and genomics platforms.

Newly generated DNA methylation data was collected from four different institutions: Henry Ford Hospital (n=103), University of Leeds (UK) (n= 8), Chinese University of Hong Kong (n=6), and Luxembourg Institute of Health (n=54). The DNA was extracted at each institution. New DNA methylation data from Henry Ford Hospital and from Chinese University of Hong Kong was generated at the University of Southern California. Briefly, the DNA was bisulfite-converted (Zymo EZ DNA methylation Kit; Zymo Research) and profiled using an Illumina Human EPIC array (EPIC). For the Luxembourg Institute of Health samples, DNA methylation data was generated by the Illumina EPIC array at the Helmholtz Zentrum München (Research Unit of Molecular Epidemiology, Institute of Epidemiology, German Research Center for Environmental Health, Neuherberg, Germany) or by the Laboratoire National de Santé (Neuropathology Unit, National Center of Pathology, Dudelange, Luxembourg). Samples from University of Leeds were profiled locally using Illumina 450K Beadchip methylation arrays. The raw DNA methylation intensity data files (IDAT) were processed with the minfi package (Fortin et al., 2017). We performed noob (Normal-exponential convolution using out-of-band probes) background correction (Triche et al., 2013) and dye bias correction using the minfi package (v 1.36.0) (Fortin et al., 2017). The DNA methylation value for each locus is presented as a beta (β) value (β = (M/(M+U)) in which M and U indicate the mean methylated and unmethylated signal intensities for each locus, respectively. β-values range from zero to one, with scores of zero indicating no DNA methylation and scores of one indicating complete DNA methylation. A detection p-value also accompanies each data point and compares the signal intensity difference between the analytical probes and a set of negative control probes on the array. Any data point with a corresponding p-value greater than 1E-4 is deemed not to be statistically significantly different from background and was thus masked as “NA”. All data files that were used in our analysis can be found at https://www.synapse.org/#!Synapse:syn17038081/wiki/585622.

RNA expression data used in this study was downloaded from the GLASS portal and the TPM data matrix was filtered for selected protein coding genes only. Next, batch effects due to the different Aliquot Batches were corrected using the COMBAT algorithm with aliquots as covariates (Leek et al., 2012).

#### Public Datasets

Public samples included in the GLASS cohort were downloaded from: TCGA/GDC (https://portal.gdc.cancer.gov), (Bai et al., 2016; Mazor et al., 2015, 2017; de Souza et al., 2018). The raw DNA methylation IDAT files were accessed and processed as described for the GLASS datasets above. Sample IDs and tissue source sites from our entire longitudinal glioma cohort are listed in Table S1.

#### Quality control

DNA methylation quality control was performed using the entire GLASS epigenetic samples to ensure the identity check of samples matched to their corresponding patient. The DNA methylation signals of probes querying high-frequency SNPs were used to calculate a pairwise agreement score across samples (Heiss and Just, 2018). Only samples that passed our pairwise agreement score cutoff were kept in the GLASS epigenetic cohort (n=357).

#### Classification of longitudinal gliomas

Longitudinal glioma samples were classified as either IDH-wildtype (Classic-like, Mesenchymal-like, LGm6) or IDH-mutant (Codel, G-CIMP-high, and G-CIMP-low) DNA methylation subtypes using the CpG methylation signatures and method previously defined by our group (Ceccarelli et al., 2016).

Our cohort was also classified into the Pan-CNS DNA methylation-based classification (Capper et al., 2018) by uploading idat files into the portal https://www.molecularneuropathology.org/mnp.

Additionally, the samples were classified into the recent transcriptomic pathway-based classification of glioblastomas (Garofano et al., 2021) using MWW-GST on the basis of the highest positive Normalized Enrichment Score (Frattini et al. 2018).

Hypermutation was defined for all recurrent tumors that had received TMZ after initial surgery and had more than 10 mutations per megabase sequenced, as described previously (Barthel et al., 2019).

#### Identification of master regulator transcription factors

Initial and recurrent glioma samples were compared using the package ELMER (Silva et al., 2019) for the following pair-wise comparison: IDHmut initial vs recurrent (20 matched pairs), IDHwt initial vs recurrent (29 pairs), and IDHmut treated vs untreated (20 matched pairs). Only samples that have both DNA methylation and gene expression available were used in these analyses.

We applied the default probe filter “distal,” which uses 158,803 probes that are >2 kbp from any transcription start site as annotated by GENCODE. ELMER version 2.14 was used with the following parameters: get.diff.meth(sig.diff) = 0, get.diff.meth(p_value) = 0.05, get.diff.meth (minSubgroupFrac) = 0.2, get.pair(Pe) = 0.1, get.pair(raw.pvalue) = 0.05, get.pair(filter.probes) = FALSE, get.pair(diff.dir)=both, get.pair(permu.size) = 100, get.pair(minSubgroupFrac) = 0.4, get.enriched.motif(lower.OR) = 1.1, get.enriched.motif(min.incidence) = 2. For each comparison, the TF subfamilies were inferred from the TFs, which had the most significant anticorrelation scores. The candidate MRTFs were next identified within the TF subfamily with FDR < 0.05 using the classification from TFClass.

#### Supervised analysis associated with treatment

Only patients with known TMZ and RT treatment status after initial surgery were included in the supervised analysis. In total, we identified 6 patients who received both TMZ and RT, 12 patients who received only TMZ, 18 patients who received RT only and 33 patients who did not receive additional treatment besides surgery. We used the Kruskal–Wallis test by ranks followed by multiple testing using the Benjamini & Hochberg (BH) method for false discovery rate (FDR) estimation (Benjamini et al., 2001) to identify differentially methylated sites between these four groups at first recurrence. To define significant CpG probes, we selected probes with FDR < 0.01 and mean DNA methylation difference > 20%.

We investigated the association of differentially methylated CpG probes (FDR < 0.01; N = 5,218 CpG probes) with gene expression from 6 untreated and 14 treated (TMZ, RT and TMZ+RT) first recurrent samples with available DNA methylation and RNA-sequencing data using ELMER and filtering by distal probes (Silva et al., 2019). Finally, we applied the function “get.enriched.motif” with only “A” motif quality score (highest confidence) to identify CpG-gene pairs enriched for the same DNA motif signature. The CpG probes and genes of interest identified by these analyses were then sent to our collaborators who are part of the GLASS-NL consortium for validation of our results.

#### DNA methylation and gene expression data from the GLASS-NL cohort

The GLASS-NL consortium has collected material from 100 IDH-mutant astrocytoma (1p19q non-codeleted) patients who underwent at least two surgical resections. Material for analysis had to be available for both resections, and the surgical interval between resections was > 6 months. Detailed clinical data, imaging, and treatment data of patients was collected within the consortium. All institutions obtained ethics approval from their institutional review boards or ethics review committees before initiation of the project. All patients provided written informed consent according to local and national guidelines.

DNA and RNA were isolated from formalin-fixed paraffin-embedded (FFPE) tumour samples as previously described (Draaisma et al., 2020). Evaluation of the area with highest tumor content was done by the pathologist (PW) on a hematoxylin and eosin stained section. Macrodissection of the marked area was then done on 10-20 10μm consecutive slides. DNA and RNA extraction was performed using the QIAamp DNA FFPE and RNeasy FFPE kit respectively (both Qiagen, Venlo, The Netherlands). DNA methylation profiling was performed with the Infinium MethylationEPIC BeadChip according to the manufacturer’s instructions making use of the Infinium FFPE DNA Restoration Kit. RNA-sequencing was done by Genomescan (Leiden, the Netherlands) and data processing, alignment and further analysis of read counts was done as described (Hoogstrate et al., 2020).

#### Deconvolution analysis

We first constructed a signature matrix from reference DNA methylation profiles of pure flow-sorted populations of cells from the literature. This signature matrix represents a set of differentially methylated CpGs selected and weighted to reflect specificity for a given cell type and is used as the basis of cell deconvolution by methylCIBERSORT. Our final signature matrix consisted of 10 cell types: CD19+ cells (B cells) (n=6), CD8+ T cells (n=6), CD56+ (natural killer cells) (n=6), and neutrophils (n=12) were from the FlowSorted.Blood.450k Bioconductor package version 1.30.0 (Reinius et al., 2012). CD4+ effector T cells (n=6) and T regs (n=4) were from (Zhang et al., 2013), accessed through the MethylCIBERSORT R package (Chakravarthy et al., 2018). Vascular endothelial cells (n=2) data was from (Moss et al., 2018). Monocyte-derived macrophage (n=4) data was from (Dekkers et al., 2019). Neuron (n=31) and glia cells (n=31) were from (Gasparoni et al., 2018). The MethylCIBERSORT R package was used to derive the DNA methylation signature for the deconvolution and the signature matrix was exported and uploaded to the CIBERSORTx portal to be deconvoluted using 1000 permutations without quantile normalization.

#### Validation of tumor cell composition

Sections of formalin-fixed, paraffin-embedded human glioma surgical samples were deparaffinized with xylene and rehydrated through graded alcohol into deionized H_2_0. Antigens were unmasked by incubation for 45 mins at 95°C in Diva Decloaker (Biocare, DV2004) using Biocare’s Decloaking chamber, and sections were stained with the antibodies listed on the table, visualized with intelliPATH FLX™ DAB Chromogen Kit (Biocare, IPK5010) and counterstained with intelliPATH™ Hematoxylin (Biocare, IPCS5006L). The antibodies are listed in the Key Resource Table.

Images of the tissue sections stained by immunohistochemistry using CD163, CD31, and CD8 antibodies were captured by an Olympus IX70 microscope and a digital camera. For quantitative analysis, we selected eight representative areas in each section. Images of the representative areas were captured at a 10X magnification. For CD163 and CD31 stainings, individual cells per area were identified by strong brown stain and counted by using ImageJ (NIH, Bethesda USA) by an algorithm to evaluate staining using hematoxylin and DAB staining specific built-in color deconvolution plug-in. For CD8 immunostaining, positive cells identified in each area by strong brown stain were manually counted. The cell counting was repeated three times. All images were analyzed in a blinded fashion. Values are presented as mean±standard deviation. Groups were compared and potential differences were identified using the non-parametric t test.

#### Statistical analysis

Data visualization and statistical analysis were performed using R version 4.1.0 software packages (www.r-project.org) and Bioconductor (Gentleman et al., 2004). Unless specified, significance was access with an FDR adjustment of less than 5%.

#### Code and data availability

Processed data for the GLASS consortium is available on Synapse (https://www.synapse.org/#!Synapse:syn21589818) and will be publicly available on November 9, 2021.

## Acknowledgements

This work is supported by the National Institutes of Health under grant numbers R01CA222146 (H.N.), R01CA222146 (I.D., L.M.P.), R01NS096236 (E.G.V.M.), R01NS117666 (E.G.V.M.), R01NS042645 (S.B.), P50CA190991-07 (M.K.); the Department of Defense grant number CA170278 (T.S.S., L.M.P., H.N.); São Paulo Research Foundation (FAPESP) grant numbers 2018/00583-0 and 2019/14928-1 (T.M.M.); Leeds Cares grant [9R11/14-11] and the Sidney Driscol Neuroscience Foundation Contribution to Brain Tumour Northwest (L.F.S.); University of Colorado Department of Neurosurgery Nervous System Biorepository (D.R.O.); FNR CORE C20/BM/14646004 (GLASS-LUX) and FNRS-Televie TETHER (S.P.N.); FNRS-Televie TETHER (A.C.H.); F.S.V. is supported by the JAX Scholar Program and a postdoctoral fellowship from The Jane Coffin Childs Memorial Fund for Medical Research; GLASS-NL was supported by the Dutch Cancer Society KWF grant number 11026 (B.Y., B.A.W., P.W., M.C.M.K., W.V., P.J.F., M.J.V.D.B., J.M.N., M.S.).

## KEY RESOURCE TABLE

| REAGENT or RESOURCE | SOURCE | IDENTIFIER |
| --- | --- | --- |
| **Software and Algorithms** |  |  |
| MethylCIBERSORT | (Chakravarthy et al., 2018) | https://doi.org/10.5281/zenodo.1298968 |
| CIBERSORTx | (Newman et al., 2019) | https://cibersortx.stanford.edu/ |
| R version 4.0.5 | (R Core Team, 2017) | https://www.R-project.org |
| minfi (v1.36.0) | (Aryee et al., 2014) | https://bioconductor.org/packages/release/bioc/html/minfi.html |
| TCGAbiolinks (v2.18.0) | (Colaprico et al., 2016) | http://bioconductor.org/packages/TCGAbiolinks/ |
| ELMER (v2.14.0) | (Silva et al., 2019) | http://bioconductor.org/packages/ELMER/ |
| DNA methylation-based classification of central nervous system tumors | (Capper et al., 2018) | https://www.molecularneuropathology.org/mnp |
| CD8 | Rabbit Monoclonal (C8/1779R) | Biocare, API3219AA |
| CD31 | Mouse Monoclonal (BC2) | Biocare, PM347 |
| CD163 | Mouse Monoclonal (10D6) | Biocare, PM353AA |
| **Deposited Data** |  |  |
| Raw DNA methylation data | This paper |  |
| TCGA data | NIH Genomic Data Commons (GDC) | https://gdc.cancer.gov/about-data/publications/PanCanStemness-2018 |
| Published longitudinal glioma cohort | (Mazor et al., 2015) | EGAS00001001255 |
| Published longitudinal glioma cohort | (Mazor et al., 2017) | EGAS00001001854 |
| Published longitudinal glioma cohort | (Bai et al., 2016) | EGAS00001001588 |
| Neuron and glia cells data | (Gasparoni et al., 2018) | GSE66351 |
| Blood cells (CD19+, CD8+ T, CD56+, neutrophils) | (Reinius et al., 2012) | GSE35069 |
| CD4+ effector T and T regs | (Zhang et al., 2013) | GSE49667 |
| Vascular endothelial cells | (Moss et al., 2018) | GSE122126 |
| Monocyte-derived macrophage | (Dekkers et al., 2019) | GSE118696 |

## Author Contributions

Conceptualization, T.M.M., T.S.S., R.G.W.V., A.I., H.N.; Methodology, T.M.M., T.S.S., H.N.;

Formal Analysis, T.M.M., T.S.S., I.D., W.V., L.G., P.J.F.;

Investigation, T.M.M., T.S.S.; Resources, T.M.M., T.S.S., F.S.V.;

Data curation, T.M.M, T.S.S, I.D., F.S.V., L.G., W.V., K.A., L.D., S.B., J.S.B., H.K.G., M.H., A.C.H., K.C.J, M.K., E.K., M.C.M.K., S.M., S.P.N., J.M.N., D.R.O., S.H.P., G.R., P.A.S.S., M.S., L.F.S., M.J.V.D.B., E.G.V.M., A.W., T.W., M.W., B.A.W., B.Y., P.W., A.L., P.J.F., L.M.P., R.G.W.V., A.I., H.N.;

Writing-Original draft, T.M.M., T.S.S., H.N., A.I.;

Writing-Review & Editing, T.M.M, T.S.S, I.D., F.S.V., L.G., W.V., K.A., L.D., S.B., J.S.B., H.K.G., M.H., A.C.H., K.C.J, M.K., E.K., M.C.M.K., S.M., S.P.N., J.M.N., D.R.O., S.H.P., G.R., P.A.S.S., M.S., L.F.S., M.J.V.D.B., E.G.V.M., A.W., T.W., M.W., B.A.W., B.Y., P.W., A.L., P.J.F., L.M.P., R.G.W.V., A.I., H.N.;

Funding acquisition, H.N., L.M.P., R.G.W.V.;

Supervision, H.N., A.I., R.G.W.V.

All co-authors and contributors discussed the results and commented on the manuscript.

