## Supplemental Figures for "The epigenetic evolution of gliomas is determined by their IDH1 mutation status and treatment regimen"

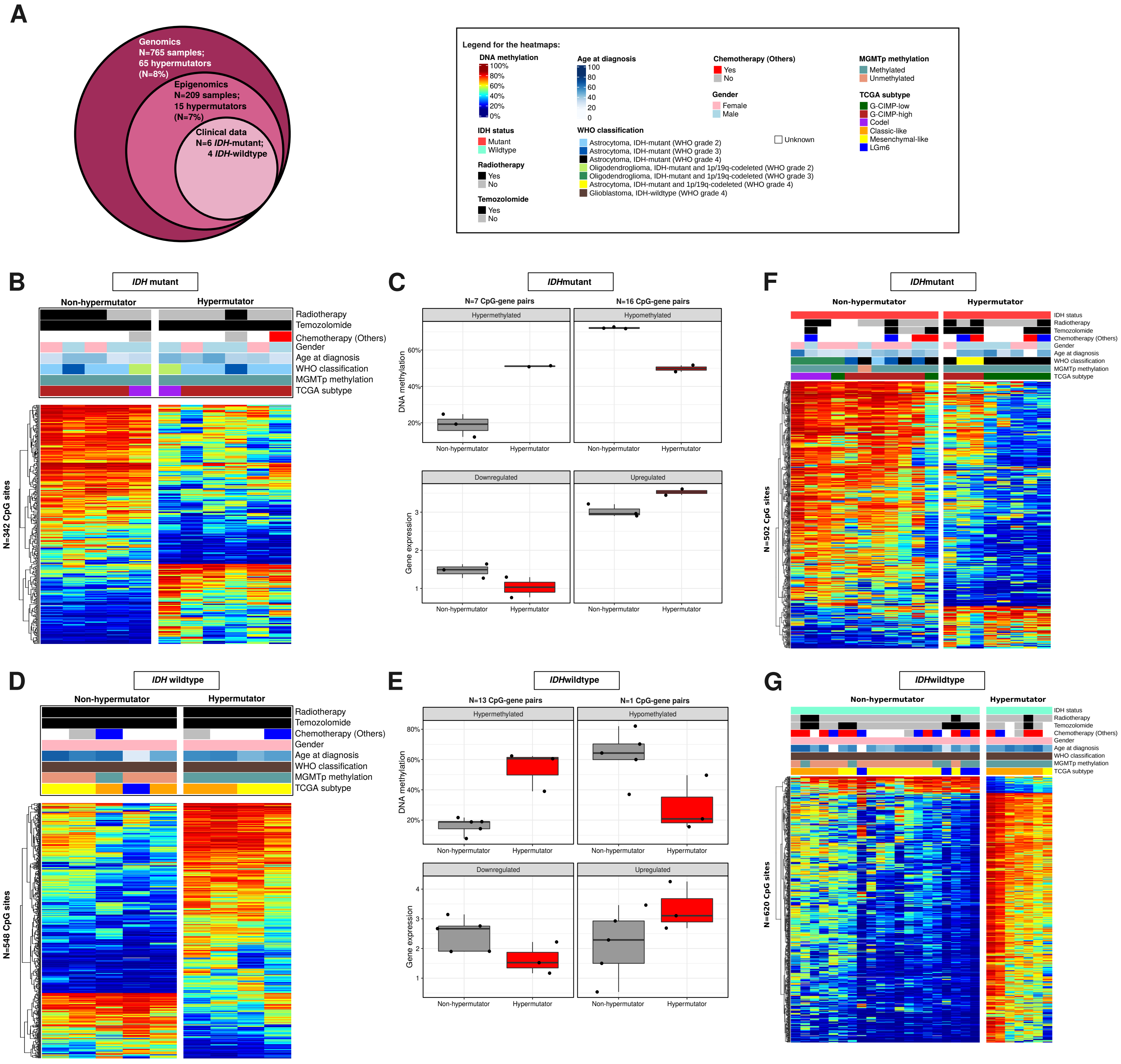


**Figure S1: DNA methylation signature to predict the hypermutator phenotype upon initial surgery divided by IDH status, related to Figure 1.** A) Venn diagram of the GLASS hypermethylator cohort based on data type: Whole-genome sequencing, DNA methylation array and clinical follow-up. B) Heatmap of DNA methylation data across IDHmut samples at initial diagnosis that were treated with TMZ, stratified by hypermutator status. Supervised hierarchical clustering analysis of 342 CpG probes that can predict the hypermutator status at the initial surgery. Column-wise represents 11 glioma samples, row-wise represents CpG probes. Samples and CpGs are ordered using hierarchical clustering method. DNA methylation beta-values range from 0 (low) to 1 (high). Additional tracks are included at the top of the heatmaps to identify each sample membership within separate cluster analysis. C) Boxplot of the CpG-gene pairs at gene promoter regions of IDHmut gliomas divided by DNA methylation (top) and gene expression (bottom) data. D) DNA methylation heatmap of 548 CpG data across IDHwt samples at initial diagnosis treated with TMZ, stratified by hypermutator status. E) Boxplot of CpG-gene pairs at gene promoter regions identified in IDHwt gliomas. F/G) Differentially methylated regions between hypermutator vs non-hypermutator recurrent samples divided by IDH status: F, IDHmut and G, IDHwt.

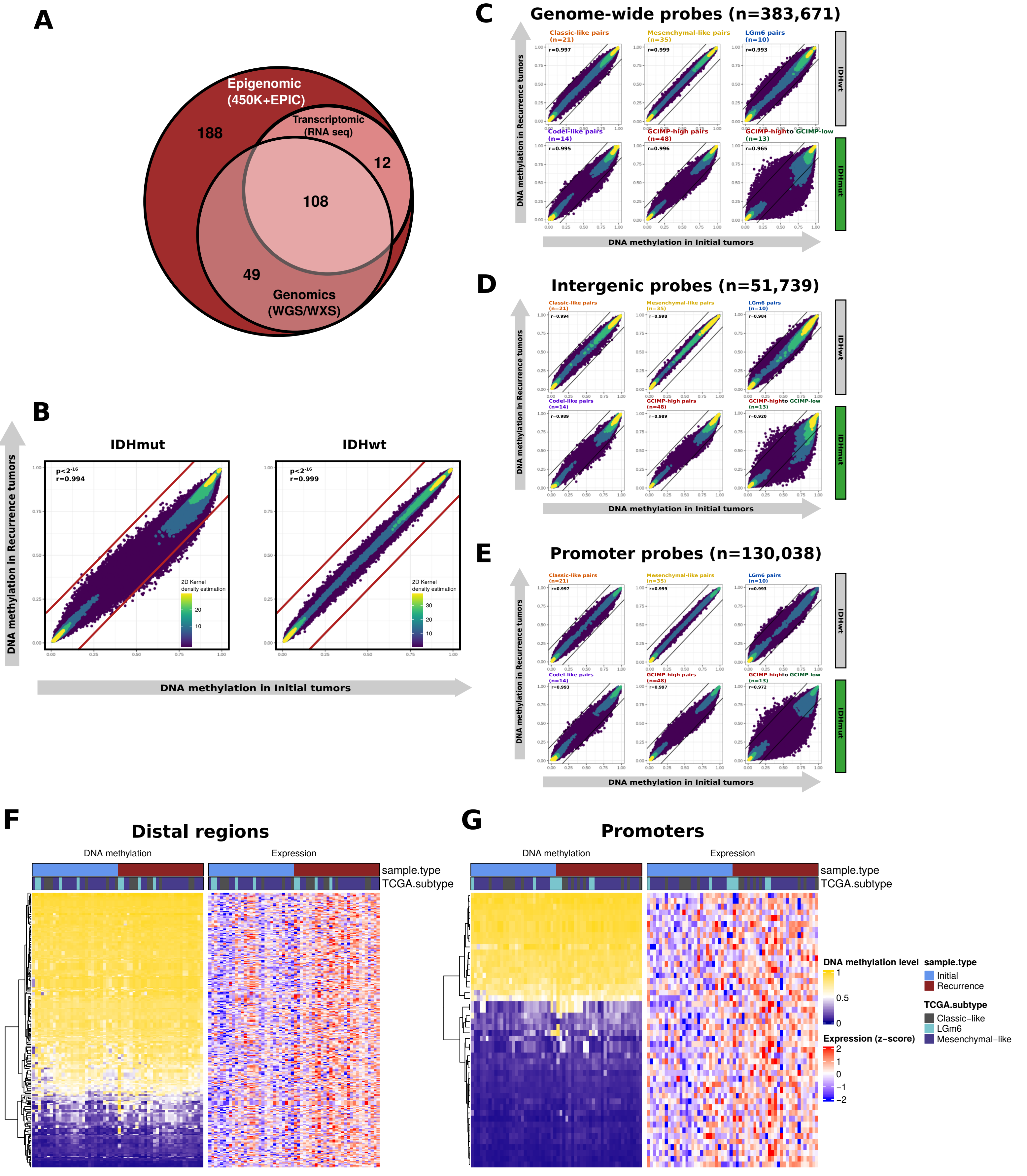


**Figure S2. Longitudinal epigenomic changes of matched initial and recurrent gliomas, related to Figure 2.** A) Venn diagram of DNA methylation samples which were profiled with genomics (WGS/WXS) and/or RNAseq. B) Overall genome-wide DNA methylation correlation between initial and recurrent tumors, stratified by IDH mutation status. C/D/E) Overall DNA methylation correlation between initial and recurrent tumors of different glioma subtypes, by CpG probe genomic location: C, genome-wide; D, probes located in intergenic regions, and E, CpG probes located in promoters of genes. F/G) Differentially methylated CpG probes defined between recurrent and initial IDHwt tumors and their associated putative genes. F, CpG probes located at distal regions and G CpG probes located at promoter regions.

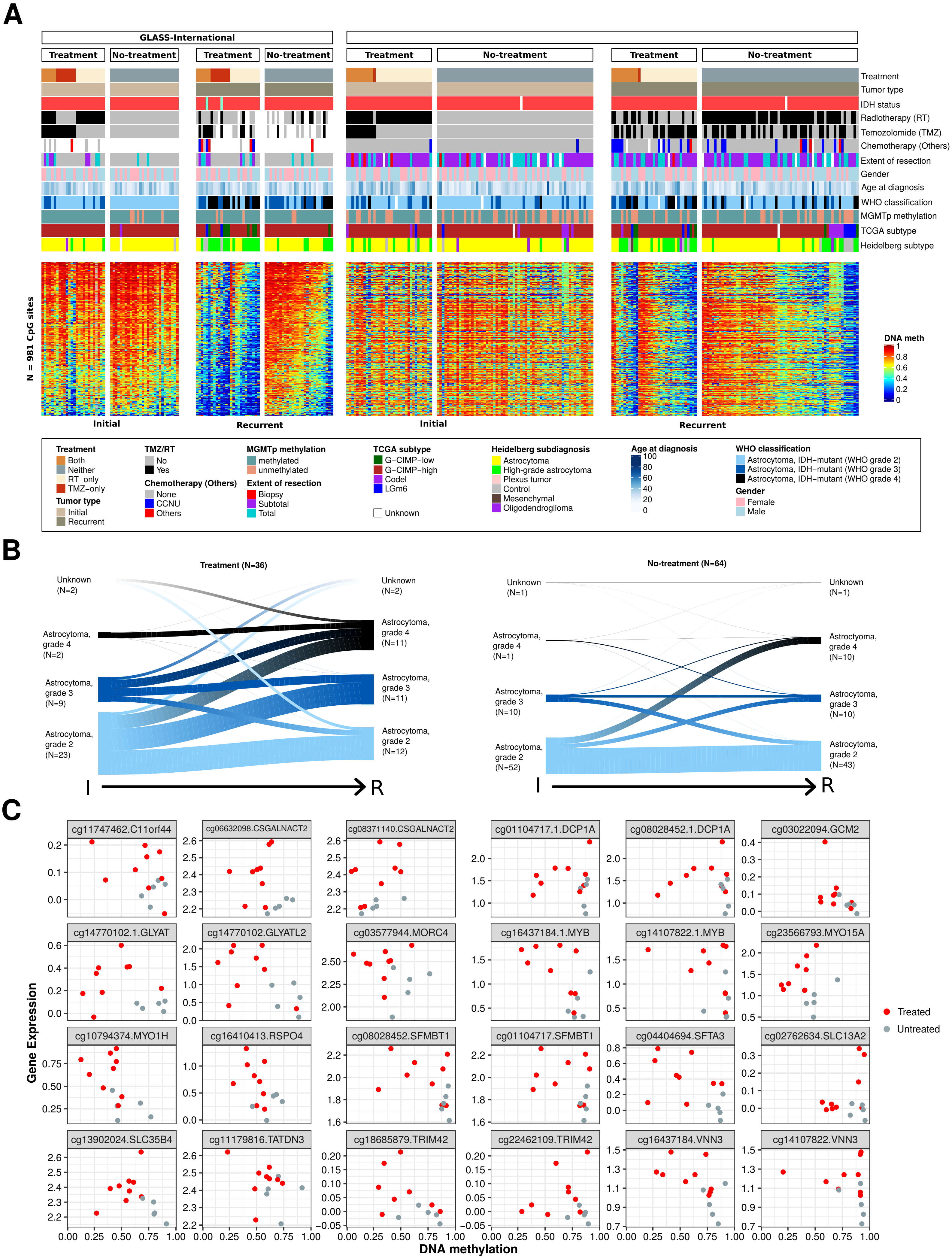


**Figure S3. DNA methylation loss is associated with malignant progression of glioma after standard treatment in the validation cohort: GLASS-NL, related to Figure 3.** A) Heatmap of DNA methylation data. Supervised analysis using astrocytoma-only samples from GLASS-International cohort identified 981 CpG probes that are associated with treatment astrocytomas IDHmut paired glioma samples. Samples from the validation cohort are also shown. Samples are stratified by cohort, initial/recurrent status and treatment status. Column-wise represents glioma samples, row-wise represents CpG probes. DNA methylation beta-values range from 0 (low) to 1 (high). Additional tracks are included at the top of the heatmaps to identify each sample membership within separate cluster analysis. B) Evolution of tumor histology (2021 WHO classification) of the validation cohort (GLASS-NL) after treatment compared to non-treated gliomas. C) Scatter plot of 24 CpG-gene pairs epigenetically regulated genes after treatment in IDHmut gliomas.

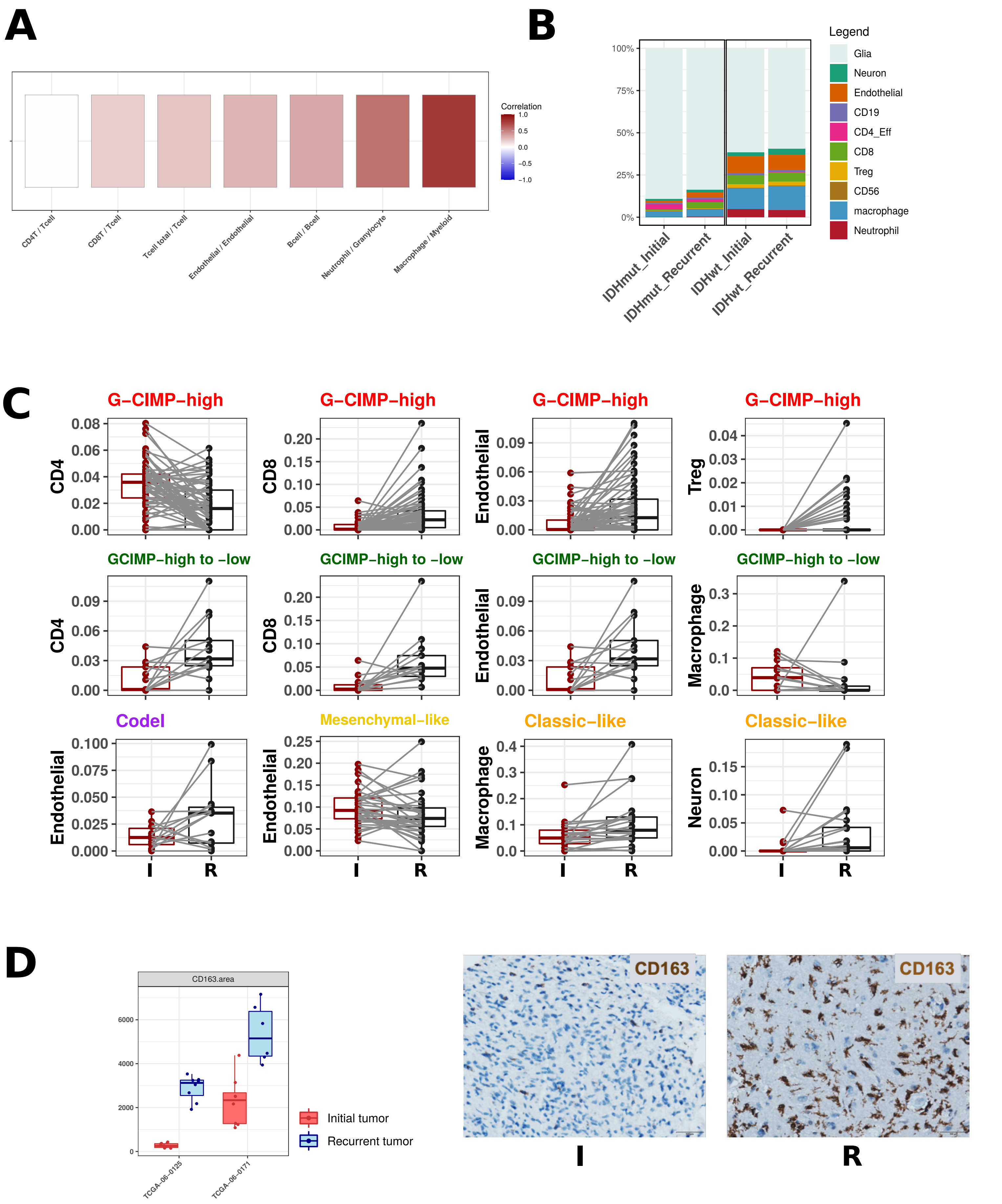


**Figure S4. Glioma subtypes present different tumor microenvironments and it changes overtime, related to Figure 4.** A) Correlation between cell composition estimated by DNA methylation and by gene expression. B) Barplots of the estimated median infiltration of specific cell types as a proportion of all cell types (range scaled from 0 to 100%) in 143 glioma tumors at initial diagnosis, divided by IDH mutation status. C) Cell type proportion (range scaled from 0 to 100%) in samples originating from the matched initial and recurrent tumors, divided by molecular subtypes. All comparisons (initial vs. recurrence, by subtype, for the specified cell population) are statistically significant ($P < 0.05$). P values calculated using a two-sided Wilcoxon rank-sum test. Matched primary and recurrent tumors are linked by the lines. D) Representative immunohistochemical staining for CD163 marker protein in two individual patients with changing levels of tumor-infiltrating immune cells
